# Differential gene expression in the interactome of the Human Dopamine transporter in the context of Parkinson’s disease

**DOI:** 10.1101/2020.10.12.336388

**Authors:** Roberto Navarro Quiroz, Ana Ligia Scott, Eric Allison Philot, Linda Atencio, Cecilia Fernandez Ponce, Gustavo Aroca Martinez, Andres Cadena Bonfanti, Lorena Gomez Escorcia, Elkin Navarro Quiroz

## Abstract

**Background:** The human dopamine transporter is the main regulator of dopamine tone and an intricate network exists to regulate the expression, conformation, and kinetics of the hDAT. hDAT dysfunction is directly related to Parkinson’s disease. The objective of this work is to evaluate the differential gene expression in the interactome of the Dopamine transporter in the context of Parkinson’s disease.

**Methods:** To do this, we evaluated hDAT interaction data in string-db, mint.bio, IntAct, reactome, hprd and BioGRID, subsequently, data was obtained from the differential gene expression of mRNA and miRNAs for this hDAT interactome in the context of PD.

**Results:** The analysis of the differential expression changes of genes of the hDAT interactome in tissues of patients with PD compared with tissues of individuals without PD, allowed to identify an expression pattern of 32 components of the hDAT interactome, of which 31 presented a negative change proportion in PD.

**Conclusions:** We found a total of 90 miRNAs that could regulate the expression of 27 components of the hDAT interactome, at the same time, 39 components of the hDAT interactome may participate in 40 metabolic pathways. Together, these findings show a systematic effect on the hDAT-mediated dopamine internalization process in patients with Parkinson’s, which would contribute to a greater susceptibility to neuronal oxidative stress in PD patients.

## Background

The global burden of Parkinson’s disease(PD) has more than doubled in the last 30 years which may be consequence of the increase population over 60 years old[1]. But, the of Parkinson’s disease continues to elude science as well as the causes of the preferential vulnerability of nigrostriatal dopaminergic neurons in PD[2].

The human dopamine transporter (hDAT) plays a key role in dopamine homeostasis in dopaminergic neurons, since it internalizes dopamine into the cell and it stores in intracellular vesicles, thus modulating the intensity and duration of signaling of dopamine [3]. The proper functioning of hDAT implies not only the formation of complexes with direct physical protein-protein interaction (PPI), as for example, structurally hDAT exhibits at its amino and carboxyl termini phosphorylation sites to multiple protein kinases[4], but it is also regulated by indirect interactions, without PPI, such as the enzymes that participate in the synthesis of dopamine,which we will refer together to as interactome.

In turn, recycling of dopamine by hDAT is especially important due to the fact that dopamine acts as antioxidant and to protect dopaminergic neurons from oxidative stress by scavenging free radicals, likewise dopamine is an important neurotransmitter that is present in high concentration in the axon terminals of dopaminergic neurons is estimated at 47 mM[5]. There is strong evidence that the redox reactions of these neurotransmitters are involved in the first steps and in the progression of neurodegenerative diseases such as Parkinson’s disease[6]. In addition, a significant reduction of dopaminergic neurons is possible to be associated with some characteristic symptoms of PD as: akinesia and tremors, conversely, abnormally high amounts of dopamine can result in hyperkinesia, altered behavior, and delusions[2, 7, 8].

Alterations in PD gene expression have been described [9, 10], both in experimental models and in humans[11–16],however, the expression of genes associated with the hDAT interactome and functional partner in Parkinson’s disease remains poorly understood [17]. A better understanding of how changes in gene expression in the hDAT interactome in PD could help clarify its role in the development of this disease and guide advances in drug discovery. For this reason this study is to evaluate the differential gene expression in the interactome of the hDAT in the context of Parkinson’s disease.

## Methods

### SLC6A3 interactome

Results from purchased with six databases of interactome (BioGRID3.5.186[18], string-db[19], mint.bio[20], IntAct[20, 21], reactome [22] and hprd[21]) was obtained using two terms: Q01959 and SLC6A3 (Sodium-dependent dopamine transporter), this in order to find proteins that had previously been related to SLC6A3.

### Enrichment analysis

It was done a GO Enrichment Analysis Powered by PANTHER[23] to relate the results coming from the interactome databases with other metabolic pathways. It was applied performed Fisher’s Exact and its correction to calculate False Discovery Rate during the analysis.

### ARNm expression Data sources

Microarrays data related to parkinson Disease were obtained from the website GEO(The Gene Expression Omnibus) (https://www.ncbi.nlm.nih.gov/geoprofiles)[24]. The following terms (‘parkinson’s disease [All Fields] AND (“CSK“OR” EPN1“OR” EPS15” OR “slc6a3” OR“MTNR1A” OR“NEDD4” OR“PARK2” OR“PICK1” OR“SNCA”OR “SNTA1“OR” STX1A“OR” TGFB1I1“OR” TUBA1B “OR"TUBB1“OR” DRD2“OR” SLC18A2 “OR“TH” OR“FLOT1 “OR”FLOT2 “OR”CDH1 “OR“DRD4”OR” DRD1“OR” TPPP“OR” RIT2“OR” GPR37 “OR“WLS”OR” Ywhae“OR” Patj“OR” Nos1“OR” Syn1“OR” HIC5“OR” RPS6KA2“OR” RACK1“[Gene Symbol] AND “Homo sapiens”[Organism]) was used in the search. The result that contained ARNm of protein associated with the interatome of the Dopamine transporter in parkinson were screened for further analysis with function GEO2R to compare two or more groups of Samples in order to identify genes that are differentially expressed across experimental conditions (False discovery rate[25]).

### Literature reviewing to miRNAs associated with parkinson Disease

Studies published before June 22, 2020 reporting the relationship between miRNAS expression and parkinson disease were retrieved from PubMed, Web of Science, EMBASE, Cochrane Central Register of Controlled Trials, Science Direct, Google Scholar, The searching terms were as follows: (“parkinson disease”[MeSH Terms] OR (“parkinson”[All Fields] AND “disease”[All Fields]) OR “parkinson disease”[All Fields] OR (“parkinson’s”[All Fields] AND “disease”[All Fields]) OR “parkinson’s disease"[All Fields]) AND (“micrornas“[MeSH Terms] OR “micrornas”[All Fields] OR “mirnas”[All Fields]). Meanwhile the literature was crossed with the database of TarBase to determine if the genes that have previously been described with a differential expression in Parkinson patients interact with some of the 34 mRNA of the proteins associated with hDAT interatom[26].

### Network constructions

Cytoscape 3.8.0 (https://cytoscape.org/) was used to build the Networks [27] and the results of miRNA analysis that previous studies had Predicted Functional Partners with mRNA of the 34 proteins that have been described as part of the hDAT interactome.

### Statistical analysis

A meta-analysis was performed to estimate the pooled effect size of ARNm expression fold 2 change and it applied a simple scaling normalization method. Study of heterogeneity was assessed using the I2 index and the heterogeneity represented by the I2 index was interpreted as modest (I2 ≤ 25%), moderate (25% < I2 ≤ 50%), substantial (50% < I2 ≤ 75%), or considerable (I2 > 75%)[28]. A fixed-effect model would be estimated when modest to moderate heterogeneity was present, and a random-effect model would be estimated when substantial to considerable heterogeneity was present. All statistical analyses were conducted using the R version 3.6.3 (2020-02-29), Package ‘meta’[29],“limma[30]” and GEO2R[31]. All analyses used two-sided tests, and p-values less than 0.05 were considered statistically significant, using the procedure of Benjamini and Hochberg. Fixed effect and random effects meta-analysis of fold change means to calculate an overall mean; inverse variance weighting is used for pooling and for random effects meta-analysis based on a refined variance estimator for the treatment estimate. which proposed a t distribution with K-2 degrees of freedom where K corresponds to the number of studies in the meta-analysis, to represent differential expression of features of SLC6A3 interactome on the x-axis we typically find the fold change and on the y-axis the −log10(p-value) (Volcano plots)(figure 2.).

### Inclusion criteria

Inclusion criteria were used for each of the data were classified according to:

a. Interactome : For each dataset were selected the protein that according to the dataset are members of the interactome of hDAT (Q 01959).
b. ARNm expression Data sources: All results of an mRNA expression experiment in a Parkinson’s context were selected from GEO and contained within information any of the components of the interatome hDAT.
c. miRNAs associated with parkinson Disease: All the results of PubMed, Web of Science, EMBASE, Cochrane Central Register of Controlled Trials, ScienceDirect and Google Scholar were selected the articles that underneath the titles and in the abstract the following terms parkinson and miRNAs and that interacted with mRNA of the components of the interatomic hDAT.

### Exclusion criteria

Exclusion criteria were used for each of the data were classified according to:

a. Interactome : All other information that according to the databases was not part of the hDAT interactome were excluded.
b. ARNm expression Data sources: All the results of an expression of mRNA that did not match in a Parkinson’s context were selected from GEO and that did not match within the information contained any of the components of the interatomic hDAT.
c. miRNAs: All the results of PubMed, Web of Science, EMBASE, Cochrane Central Register of Controlled Trials, ScienceDirect and Google Scholar were excluded the articles that not had in the titles and in the abstract the following terms parkinson and miRNAs and that interacted with mRNA of the components of the interatomic hDAT.

## Results

### hDAT interactors

We found a total of 39 interactor proteins for hDAT (slc6a3 UniProtKB / Swiss-Prot protein ID Q01959) in the database like: BioGRID [18]; string-db [19]; mint.bio [32]; hprd [21], Reactome [33], and IntAct [20] (figure 1).

**Figure 1.**
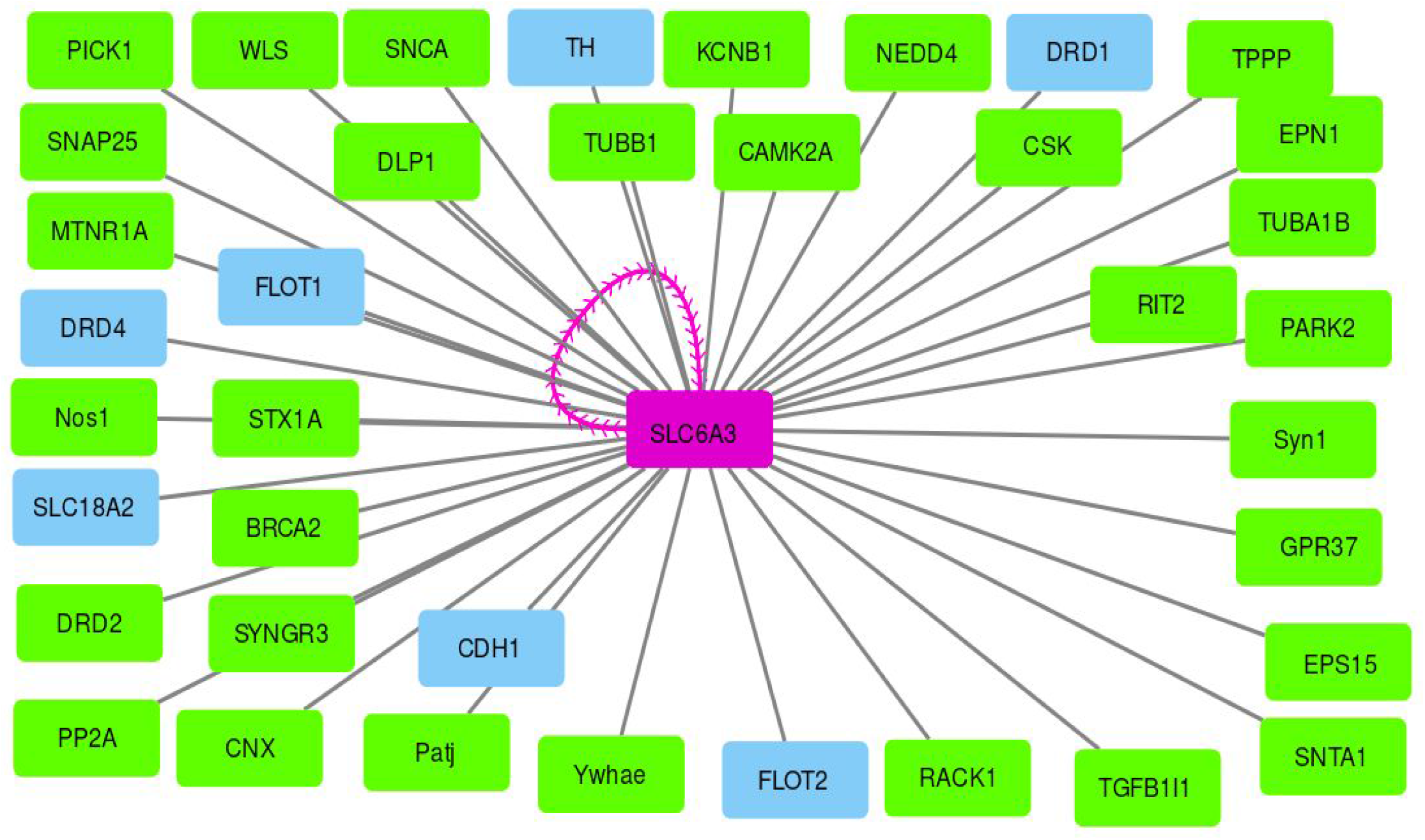
hDAT interactome. The magenta rectangle shows the SLC6A3 and the magenta line reports that there is dimer formation of SLC6A3. The green rectangles show direct protein-protein interactions and the blue rectangles are indirect interactions as functional partners.

**Figure 2.**
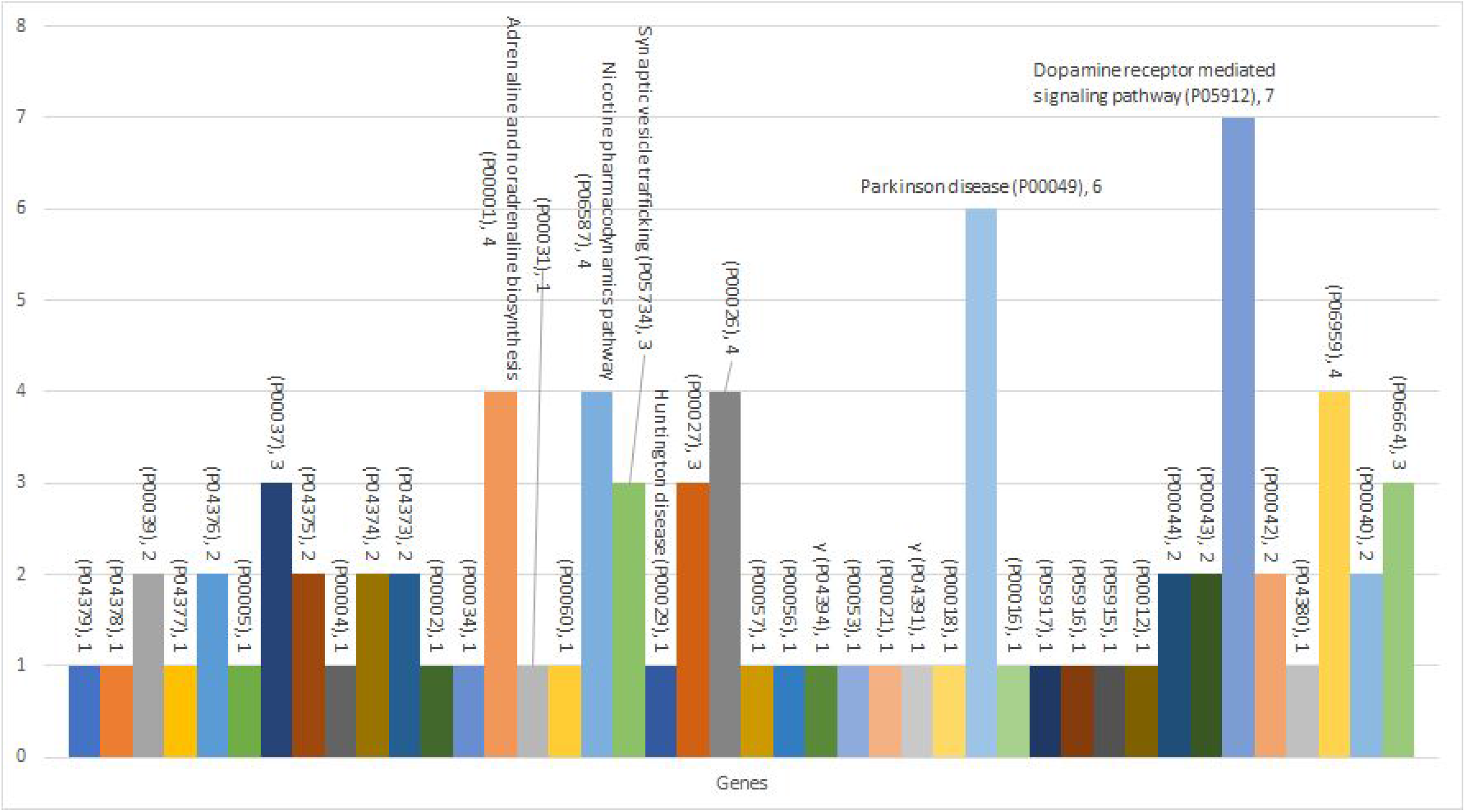
GO metabolic pathways associated with the hDAT interactome; the para labels correspond to Pathway Accession associated with neurodegenerative diseases and the metabolic pathways in which hDAT interatome has the greatest contributions

### GO: “pathway” enrichment analysis (Panther)

An Independent enrichment analysis was carried out using Panther[23] and using as search term each of the 39 components of the interactome, result in the following 40 “GO pathway”: Beta3 adrenergic receptor signaling pathway (P04379), Beta2 adrenergic receptor signaling pathway (P04378),Metabotropic glutamate receptor group III pathway (P00039),Beta1 adrenergic receptor signaling pathway (P04377), 5HT4 type receptor mediated signaling pathway (P04376),Angiogenesis (P00005), Ionotropic glutamate receptor pathway (P00037), 5HT3 type receptor mediated signaling pathway (P04375), Alzheimer disease-presenilin pathway (P00004), 5HT2 type receptor mediated signaling pathway (P04374), 5HT1 type receptor mediated signaling pathway (P04373), Alpha adrenergic receptor signaling pathway (P00002), Integrin signalling pathway (P00034), Adrenaline and noradrenaline biosynthesis (P00001), Inflammation mediated by chemokine and cytokine signaling pathway (P00031), Ubiquitin proteasome pathway (P00060), Nicotine pharmacodynamics pathway (P06587), Synaptic vesicle trafficking (P05734), Huntington disease (P00029), Heterotrimeric G-protein signaling pathway-Gq alpha and Go alpha mediated pathway (P00027), Heterotrimeric G-protein signaling pathway-Gi alpha and Gs alpha mediated pathway (P00026), Wnt signaling pathway (P00057), VEGF signaling pathway (P00056), Thyrotropin-releasing hormone receptor signaling pathway (P04394), T cell activation (P00053), FGF signaling pathway (P00021), Oxytocin receptor mediated signaling pathway (P04391), EGF receptor signaling pathway (P00018), Parkinson disease (P00049), Cytoskeletal regulation by Rho GTPase (P00016), Opioid proopiomelanocortin pathway (P05917), Opioid prodynorphin pathway (P05916), Opioid proenkephalin pathway (P05915), Cadherin signaling pathway (P00012), Nicotinic acetylcholine receptor signaling pathway (P00044), Muscarinic acetylcholine receptor 2 and 4 signaling pathway (P00043), Dopamine receptor mediated signaling pathway (P05912), Muscarinic acetylcholine receptor 1 and 3 signaling pathway (P00042), Cortocotropin releasing factor receptor signaling pathway (P04380), CCKR signaling map (P06959), Metabotropic glutamate receptor group II pathway (P00040) and Gonadotropin-releasing hormone receptor pathway (P06664).

### Transcriptional studies

A set of five transcriptomic studies were identified in GEO datasets, which met the inclusion criteria, were identified, out of a total of 55,393 obtained, these datasets consisted of four samples Brain Substantia nigra tissue, brain(inferior Olivary nucleus) and brain (Dorsal motor Nucleus of the vagus), a total of 107 parkinson patients and 96 controls. details of the microarray studies included are summarized in Table 1.

**Table 1.**
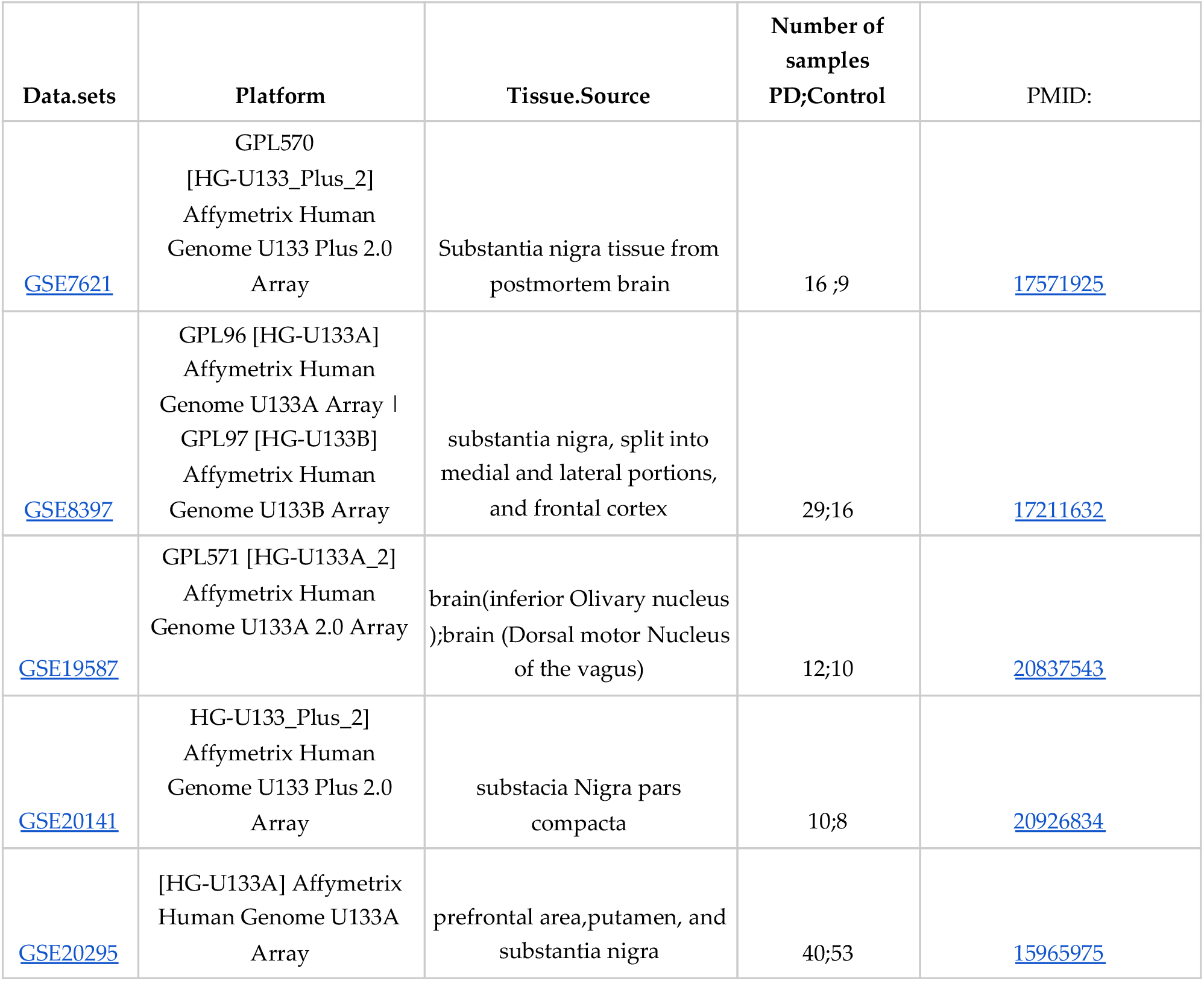
Characteristics of of the microarray studies included in this meta-analysis

| Data.sets | Platform | Tissue.Source | Number of samples<br>PD;Control | PMID: |
| --- | --- | --- | --- | --- |
| GSE7621 | GPL570<br>[HG-U133_Plus_2]<br>Affymetrix Human<br>Genome U133 Plus 2.0<br>Array | Substantia nigra tissue from<br>postmortem brain | 16 ;9 | 17571925 |
| GSE8397 | GPL96 [HG-U133A]<br>Affymetrix Human<br>Genome U133A Array \|<br>GPL97 [HG-U133B]<br>Affymetrix Human<br>Genome U133B Array | substantia nigra, split into<br>medial and lateral portions,<br>and frontal cortex | 29;16 | 17211632 |
| GSE19587 | GPL571 [HG-U133A_2]<br>Affymetrix Human<br>Genome U133A 2.0 Array | brain(inferior Olivary nucleus<br>);brain (Dorsal motor Nucleus<br>of the vagus) | 12;10 | 20837543 |
| GSE20141 | HG-U133_Plus_2]<br>Affymetrix Human<br>Genome U133 Plus 2.0<br>Array | substacia Nigra pars<br>compacta | 10;8 | 20926834 |
| GSE20295 | [HG-U133A] Affymetrix<br>Human Genome U133A<br>Array | prefrontal area,putamen, and<br>substantia nigra | 40;53 | 15965975 |

### Analysis of Gene Expression in PD

The combined estimate of the proportion of the change in 5 transcriptomic studies in the context of PD showed 32 mRNAs with altered expression with respect to the controls (Figure 4), in which we also evaluated the effect of the sample size in each of the studies (to estimate the pooled effect size of ARNm expression log2 fold change).

On the SLC6A3 interactome was possible seen differential expression to follow pool genes RIT2, SNCA and PCNX1 these downregulation, for its part, nine genes was seen upregulation (figura.3),This corresponds to the GSE20141 data set.

**Figure 3.**
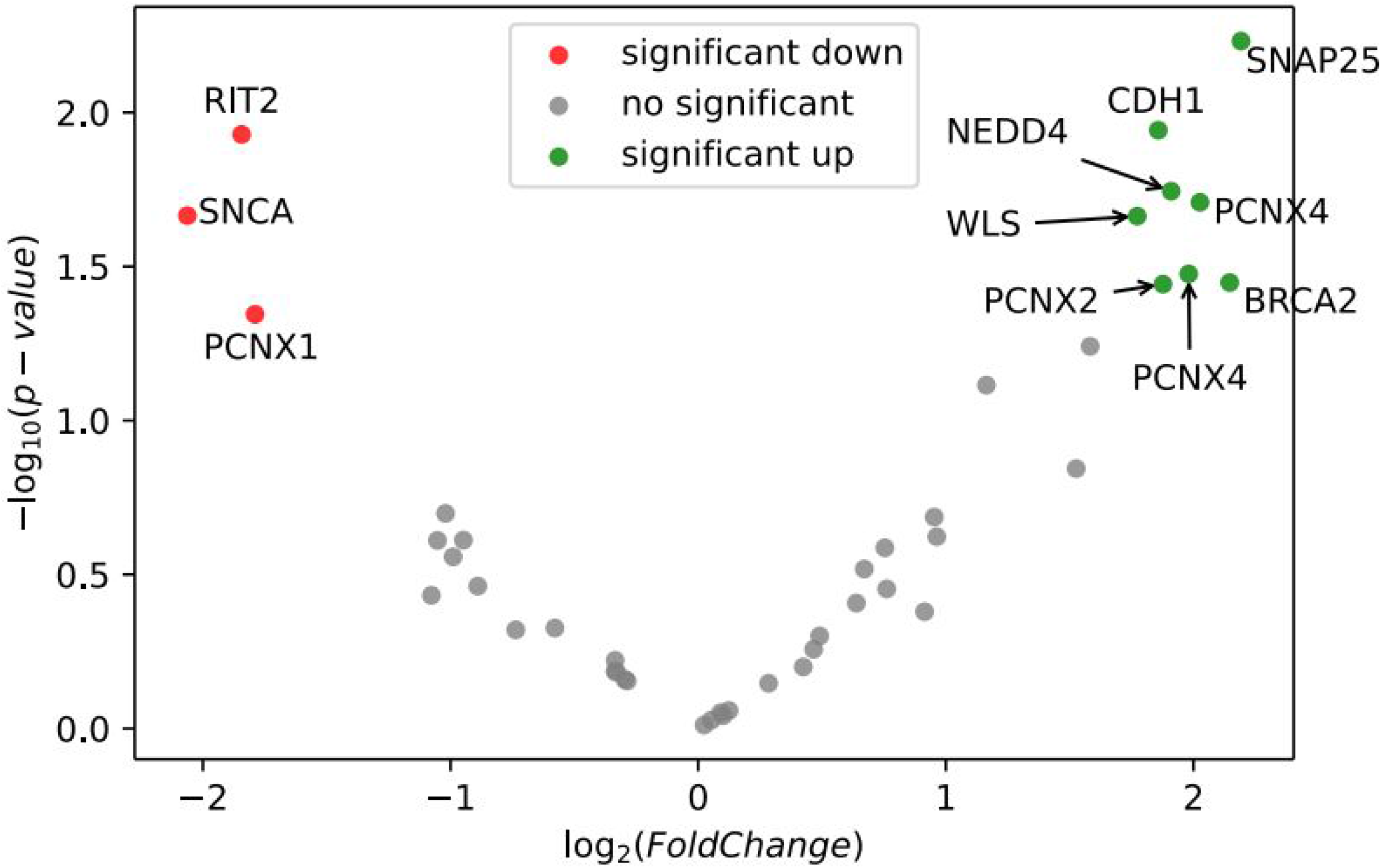
volcano plot of GSE20141 on the x-axis we typically find the fold change and on the y-axis the −log10(p-value), in red downregulation and green uprelation genes ofThe SLC6A3 interactome.

**Figure 4.**
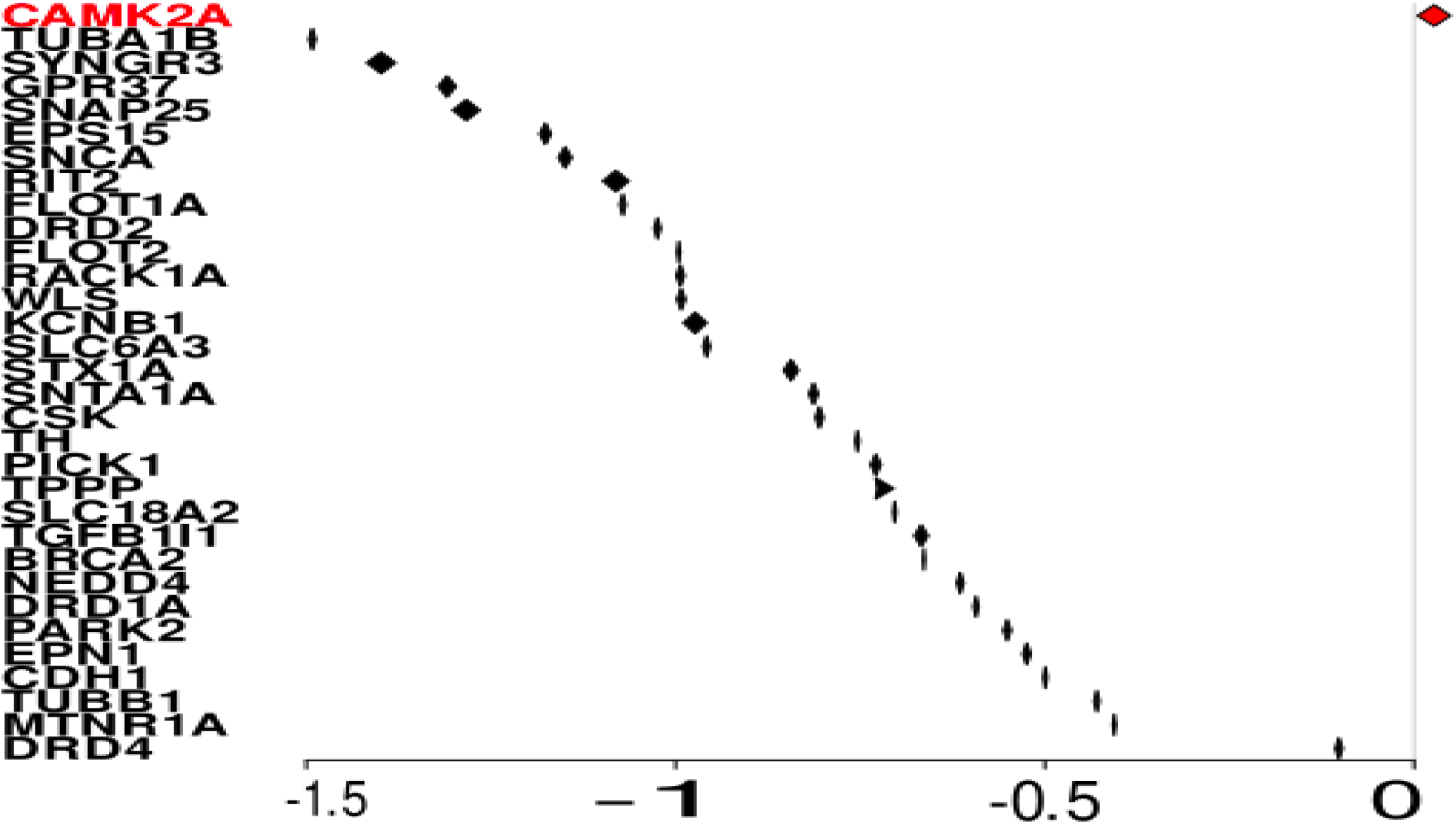
Forest plot of nine microarrays assessing 32 mRNA expression in PD. PD and normal groups. The x-axis is the estimated pooled effect size of ARNm expression fold 2 change; the size of the rhombus is associated with the range of 95% confidence intervals. and Y axis each of the 32 symbols of the genes in the evaluation of mRNA expression in PD.

The random-effect model reported, with a pooled, in this study, the random-effect model range was determined to be between 79.4% to 100%. So that the level of heterogeneity defined with significance for all mARN expressions, the heterogeneity of studies is represented by a statistical parameter representing the inter-study variation and estimating and hence the overall population treatment effect.

### The miRNAs Associated With the PD-Associated that regulate the expression of hDAT interactome mRNA

We found that 90 miRNAs with altered expression in PD had a total of 211 interactions with 27 mRNAs of the hDAT interactome. EPS15 was the mRNA of the hDAT interactome that exhibited a greater number of interactions with 24 miRNAs aberrantly expressed in PD (Figure 5).

**Figure 5.**
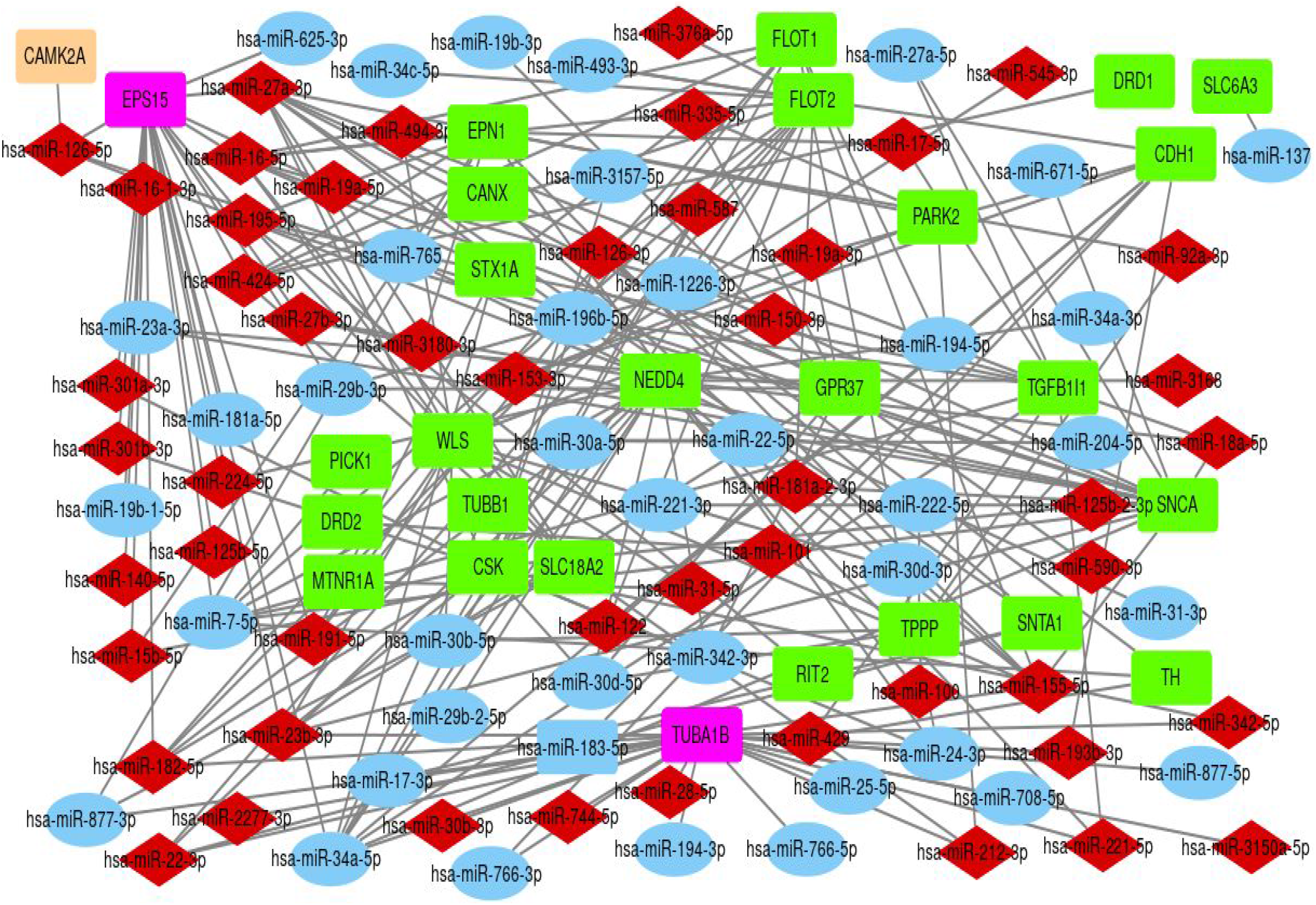
Interaction network between miRNAs and hDAT interactome mRNA in a PD context.(in red rhombus are miRNAs upregulated, light blue oval are miRNAs downregulated, violet rectangles are (TUBA1B and EPS15), pink rectangle is CAMK2A and green rectangles are mRNAs).

## Discussion

Several studies have validated the relationship between the loss of the dopamine, the posture and movement disorders characteristic of PD[34–37]. The activity of hDAT is critical in dopaminergic signaling pathways. The protein hDAT is a secondary active transport that uses ion gradients between the synaptic cleft and the presynaptic neuron to drive dopamine internalization[38]. This re-entry of dopamine within the nerve cell terminates neurotransmission and allows dopamine to be recycled for subsequent release[39]. A wide variety of proteins have been described that interact with DAT[40, 41]. These hDAT interactors are involved in a variety of actions and are key to the proper functioning of this transporter[42]. This hDAT interactome represents a complex dynamic system that is not yet fully understood. In turn, alterations in the expression of genes and of regulators of their expression such as miRNAs in PD have been shown. A greater understanding of this interactome, and of the changes in the expression of its genes in PD[15, 43], will help to understand the effects of drugs and disorders in the dopamine systems as a whole.

In this study we identified 39 interactome proteins for hDAT using UniProtKB / Swiss-Prot ID Q01959 for slc6a3 in protein interaction repositories, proteomic databases, and metabolic and signaling molecule databases (BioGRID, string-db, mint. bio, hprd, IntAct and Reactome). We show 7 Functional Partners (TH, DRD1, FLOT2, CDH1, SLC18A2, DRD4, FLOT1) and 32 heterologous direct interactors(figure 1). Subsequently, we found 5 studies of differential expression of genes by microarray of brain tissues of patients with PD compared with the same tissues of individuals without PD, where we found information on the differential expression of 32 of the 39 interactors described here for hDAT. Of these 31 mRNAs presented a combined estimate of the proportion of the negative change in PD with respect to the controls, including SLC6A3. Therefore, in general terms, an alteration in the dopamine internalization process by hDAT would be expected, which could be directly associated with a lower production of dopamine, due to the decrease that we also find in TH mRNA, which codes for tyrosine hydroxylase, catalyzes the formation of L-dihydroxyphenylalanine (L-DOPA)[44, 45], an enzyme that limits the speed in the biosynthesis of dopamine; In addition to this, the alterations found in the SNCA mRNA that encodes α-synuclein, which affects the vesicular transport and storage of dopamine, by altering the interaction with proteins such as tubulin (TUBA1B), which also had a decreased expression in its mRNA. Stored dopamine contributes to the regulation of oxidative stress in neurons, therefore this process would also be affected by the changes caused in the internalization, transport and storage of dopamine in the context of PD. Regarding the identification of SNCA in other studies in which a transcriptome meta-analysis is carried out in PD and that present some segments immediately adjacent to the segment of chromosome 4 (4q21.1) that comprises the SNCA gene (PARK1 / PARK4), which encodes alpha-synuclein [32, 43]. Furthermore, a study evaluating where proteomics (12 PD, 12 controls) and RNA sequencing transcriptomics (29 PD, 44 controls) shows that ten of the genes or proteins involved were co-localized at GWAS loci and that SNCA was stronger in proteomics than in RNA sequencing analysis [18] which is consistent with the results of the SNCA transcriptomics meta-analysis of this study.

In the same way, previous studies integrating proteomics and RNA transcriptomics in Parkinson disease[14], describe results comparable to those entered into the study for 19 mRNAs belonging to the hDAT interactome. The comparisons for each of the 19 mRNAs than those previously identified, 8 had a similar pattern of expression to those shown in this study (SNTA1, CSK, EPS15, PARK2, PICK1, DRD2, SLC18A2 and TH). Each comparison started with the result log2FoldChange PD vs. Control of its results[14] with the result estimate of the pooled effect size of ARNm expression log2 fold change of this study. However, it is pertinent to mention that the methodology used in this study, it was not possible to establish an expression pattern for Ywhae, Patj, Nos1, Syn1, CNX, DLP1 and PP2A.

In relations with the TUBA1B expresión, we observe that TUBA1B presented the lowest value estimate the pooled effect size of log2 fold change (−1.4919 [95%-CI=−1.4964; −1.4873]) to 5 transcriptomic studies in the context of PD, According to Independent Enrichment Analysis of gene sets the TUBA1B is a component of the Gonadotropin releasing hormone receptor pathway (P06664) as well as NOS1 and DRD2, The effect of Gonadotropin releasing hormone receptor pathway in PD is a controversial topic[46]; however, Gonadotropin releasing hormone receptor pathway the has been demonstrated participate in the process of Neuronal Survival [47, 48] that in parkinson’s context where is observed neuron decline subependymal zone, subgranular zone and the midbrain, It is possible to draw a picture in which alteration in the Neuronal Survival mechanism can contribute to the decrease of neurons in these areas, in addition to the TUBA1B downrelation, we propose that hsa-miR-23b-3p, hsa-miR-2277-3p, hsa-miR-342-5p, hsa-miR-155-5p, hsa-miR-30b-3p, hsa-miR-193b-3p, hsa-miR-221-5p, hsa-miR-744-5p, hsa-miR-126-5p, hsa-miR-28-5p, hsa-miR-182-5p, hsa-miR-212-3p and hsa-miR-3150a-5p (figure.5) found upregulated in parkinson patient [49] and experimentally shown to target TUBA1B[50], which the TUBA1B downrelation may have a key role in regulation of dopaminergic neurons Survival in parkinson’s context[51, 52].

Regarding the to miRNA – target interactions (MTIs) in interatome hDAT and are deregulated in PD, in a previous study of Parkinson’s patients that aimed to the Identification of Novel Biomarkers in PD was carried out as follows approach: i)experimentally supported databases with experimentally verified and ii) different prediction algorithms[13], which allows obtaining the following resulted miRNAs : miR-126-5p, hsa-miR-29b-3p, and hsa-miR-301a-3p which are also described in the present study about this miRNAs and interaction with mRNA previously described that miR-126-5p interaction with CAMK2A [53], EPS15 [53, 54], NEDD4 and SNCA[56], the hsa-miR-29b-3p interaction with EPS15 [57], CDH1 [58], GPR37[59] and the hsa-miR-301a-3p interaction with CSK[60] and EPS15[61] which are also part of hDAT interactome, which although we found miRNAs that interacted with multiple mRNAs of hDAT interactome proteins, which presented altered expression in PD vs controls, we did not find any type of relationship either in the number of miRNAs per target of the interactome, or in the combined estimate of rate of miRNA change and the combined estimate of rate of change in hDAT interactome mRNAs.

For future studies, it is possible to take into account expression and that in which regions it interacts with transcriptome mRNA, since this information may allow us to better understand the regulation of miRNAs in the hDAT interatome expression, for its part, it is advisable to carry out a study of the protein of the interactome. which can bring important information to understand the pathogenesis of parkinson and can lead to identify new markers of the disease.

## Conclusions

The analysis of the differential expression changes of genes of the hDAT interactome in tissues of patients with PD compared with tissues of individuals without PD, allowed to identify an expression pattern of 32 components of the hDAT interactome, of which 31 presented a negative change proportion in PD. We found a total of 90 miRNAs that could regulate the expression of 27 components of the hDAT interactome, at the same time, 39 components of the hDAT interactome may participate in 40 metabolic pathways. Together, these findings show a systematic effect on the hDAT-mediated dopamine internalization process in patients with Parkinson’s, which would contribute to a greater susceptibility to neuronal oxidative stress in PD patients.

## List of abbreviations

CAMK2A: CALCIUM/CALMODULIN-DEPENDENT PROTEIN KINASE II-ALPHA; CAMK2A
CSK: CYTOPLASMIC TYROSINE KINASE
EPN1: EPS-15-Interacting Protein 1
EPS15: Epsin-1
MTNR1: MELATONIN RECEPTOR 1A
NEDD4: NEURAL PRECURSOR CELL EXPRESSED, DEVELOPMENTALLY DOWNREGULATED 4
GEO: the Genetic Expression Omnibus PARK2 E3 ubiquitin-protein ligase parkin
PICK1: PRKCA-binding protein
hDAT: The human dopamine transporter
TPPP: TUBULIN POLYMERIZATION-PROMOTING PROTEIN
SLC6A3: SOLUTE CARRIER FAMILY 6 (NEUROTRANSMITTER TRANSPORTER, DOPAMINE), MEMBER 3

## Declarations

### Ethics approval and consent to participate

Not applicable

### Consent for publication

Not applicable

### Availability of data and materials

The datasets generated and/or analysed during the current study are available in the https://www.ncbi.nlm.nih.gov/geo/ and https://carolina.imis.athena-innovation.gr/diana_tools/web/index.php?r=tarbasev8%2Findex

The datasets used and/or analysed during the current study are available from the

##### WEBSIDE

https://www.ncbi.nlm.nih.gov/geo/query/acc.cgi?acc=GSE7621

https://www.ncbi.nlm.nih.gov/sites/GDSbrowser?acc=GDS4401

https://www.ncbi.nlm.nih.gov/geo/query/acc.cgi?acc=GSE8397

https://www.ncbi.nlm.nih.gov/geo/query/acc.cgi?acc=GSE19587

https://www.ncbi.nlm.nih.gov/geo/geo2r/?acc=GSE20141

https://www.ncbi.nlm.nih.gov/geo/query/acc.cgi?acc=GSE20295

